# Combining accurate tumour genome simulation with crowd-sourcing to benchmark somatic structural variant detection

**DOI:** 10.1101/224733

**Authors:** Anna Y. Lee, Adam D. Ewing, Kyle Ellrott, Yin Hu, Kathleen E. Houlahan, J.Christopher Bare, Shadrielle Melijah G. Espiritu, Vincent Huang, Kristen Dang, Zechen Chong, Cristian Caloian, Takafumi N. Yamaguchi, ICGC-TCGA DREAM Somatic Mutation Calling Challenge Participants, Michael R. Kellen, Ken Chen, Thea C. Norman, Stephen H. Friend, Justin Guinney, Gustavo Stolovitzky, David Haussler, Adam A. Margolin, Joshua M. Stuart, Paul C. Boutros

**Affiliations:** Ontario Institute for Cancer Research; Toronto, Ontario, Canada; Department of Biomolecular Engineering; University of California, Santa Cruz; Santa Cruz, CA, USA; Mater Research Institute; University of Queensland; Woolloongabba, QLD, Australia; Computational Biology Program; Oregon Health & Science University; Portland, OR, USA; Sage Bionetworks; Seattle, WA, USA; Department of Bioinformatics and Computational Biology; University of Texas MD Anderson Cancer Center; Houston, TX, USA; Department of Genetics; University of Alabama at Birmingham; Birmingham, AL, USA; Informatics Institute; University of Alabama at Birmingham; Birmingham, AL, USA; IBM Computational Biology Center; T.J.Watson Research Center; Yorktown Heights, NY, USA; Department of Medical Biophysics; Uni 23 versity of Toronto; Toronto, Ontario, Canada; Department of Pharmacology & Toxicology; University of Toronto; Toronto, Ontario, Canada

**Keywords:** somatic mutations, simulation, structural variants, benchmarking, cancer genomics, whole genome sequencing, crowd-sourcing

## Abstract

**Background:** The phenotypes of cancer cells are driven in part by somatic structural variants. Structural variants can initiate tumors, enhance their aggressiveness and provide unique therapeutic opportunities. Whole-genome sequencing of tumors can allow exhaustive identification of the specific structural variants present in an individual cancer, facilitating both clinical diagnostics and the discovery of novel mutagenic mechanisms. A plethora of somatic structural variant detection algorithms have been created to enable these discoveries, however there are no systematic benchmarks of them. Rigorous performance evaluation of somatic structural variant detection methods has been challenged by the lack of gold-standards, extensive resource requirements and difficulties arising from the need to share personal genomic information.

**Results:** To facilitate structural variant detection algorithm evaluations, we create a robust simulation framework for somatic structural variants by extending the BAMSurgeon algorithm. We then organize and enable a crowd-sourced benchmarking within the ICGC-TCGA DREAM Somatic Mutation Calling Challenge (SMC-DNA). We report here the results of structural variant benchmarking on three different tumors, comprising 204 submissions from 15 teams. In addition to ranking methods, we identify characteristic error-profiles of individual algorithms and general trends across them. Surprisingly, we find that ensembles of analysis pipelines do not always outperform the best individual method, indicating a need for new ways to aggregate somatic structural variant detection approaches.

**Conclusions:** The synthetic tumors and somatic structural variant detection leaderboards remain available as a community benchmarking resource, and BAMSurgeon is available at https://github.com/adamewing/bamsurgeon.

## Background

Somatic structural variants (SVs) are mutations that arise in tumours involving rearrangements, duplications or deletions of large segments of DNA. SVs are often defined to be events larger than 100 bp in size, although with significant variability in this definition. Somatic SVs are critical in driving and regulating tumour biology. They can initiate tumours [1,2] and because they are unique to the cancer, can serve as highly-selective avenues for therapeutic intervention [3]. The overall mutation load of somatic SVs serves as a proxy for genomic instability, and can robustly predict tumour aggressiveness in multiple tumour types [4,5].

While somatic SVs that alter copy-number can be detected using microarray assays, the resolution of such studies is limited, and many other important types of SVs cannot be detected. As a result, high-throughput DNA sequencing is now a standard approach for detecting SVs in cancer genomes. Although RNA-based assays are useful for detecting SVs that alter protein-structure, DNA-based assays are required for most others. As a result, a broad range of algorithms has been developed to detect SVs from short-read sequencing data using read depth analysis, split read (*i.e.* a read that maps to multiple different parts of the reference sequence) alignment, paired end mapping and de novo assembly techniques [6–9]. However, the accuracy of existing methods is poorly described. There are no comprehensive benchmarks of somatic SV detection approaches. Most comparison results are reported by the developers of newly published methods. These developer-run benchmarks are potentially subject to several types of selection biases. For example, the developers of one tool may be experts in parameterizing and tuning it, but may lack the same skill in tuning methods developed by others. Further, evaluating the accuracy of somatic SV detection is more challenging than evaluating the accuracy of somatic single nucleotide variant (SNV) detection as validation data is more difficult to generate for SVs. Even the metrics of measuring accuracy are not agreed upon, with no community-accepted standards on how SV prediction accuracy should be assessed, especially when predictions are close to, but not exactly at, the actual sequence breakpoints. As a result, there are no robust estimates of the false positive and false negative rates of somatic SV prediction tools on tumours of different characteristics.

To fill this gap, we created an open challenge-based assessment of somatic SV prediction tools as part of the ICGC-TCGA DREAM Somatic Mutation Calling Challenge (the Challenge). The lack of fully-characterized tumour genomes for building gold standard sets of SVs motivated our simulation approach. Specifically, we first extended BAMSurgeon [10], a tool for adding simulated mutations to existing reads, to generate somatic SVs. This approach is advantageous because it permits flexibility with the added mutations while also capturing sequencing technology biases through the use of existing reads. We created and distributed *in silico* tumours (IS1-IS3), on which 204 submissions were made by 15 teams.

## Results

### Simulation of SVs with BAMSurgeon

In addition to point mutations [SNVs and short insertions or deletions (INDELs)], BAMSurgeon is capable of creating simple SVs through read selection, local sequence assembly, manipulation of assembled contigs, and simulation of sequence coverage over the altered contigs (Fig. 1a, Additional file 1: Figure S1). This, combined with careful tracking of read depth, yields approximations of SVs including insertions, deletions, duplication, and inversions into pre-existing backgrounds of real sequence data. Here we present results based on simulations of those SV types. Subsequent to the Challenge, BAMSurgeon was extended to support translocations and more complex rearrangements. The BAMSurgeon manual, available online, contains a full description of input formatting and available parameters. The input regions define where local assembly will be attempted *via* Velvet [11]. For each region, the largest assembled contig is selected and re-aligned to the reference genome using Exonerate [12]. The contig is then trimmed to the length of its longest contiguous alignment and the alignment is used to accurately track breakpoint locations within the contig in terms of reference coordinate space. The location and identity of reads from the original BAM file in the assembled contig are tracked *via* parsing of the AMOS [13] file output by Velvet [14], which also enables tracking of reads included or excluded after contig trimming. If a suitable contig (*i.e.* sufficiently long, with a sufficiently low number of discordant read pairs) is not available for a given input region, no mutation is made for that region. For each segment where contig assembly succeeds, the contig is rearranged according to the user specification (*e.g.* insertion, deletion, duplication, or inversion of sequence). Then paired reads are simulated from the rearranged contig using wgsim [15], with specific parameters controllable by the user. Because reads are simulated using the rearranged contig, breakpoint-spanning reads and reads that will be discordant versus the reference genome assembly will be created. The number of reads simulated (final coverage, *C_f_*) depends on the original coverage *C_o_*, the difference in length between the original contig *L_o_* and the rearranged contig *L_f_*, **and** a user-specified parameter controlling variant allele fraction (VAF). Thus, *C_f_* = VAF\**C_o_*\*(*L_f_*/*L_o_*). Duplications and insertions result in larger contigs and require new reads to be **added** to the final BAM, and deletions yielding a smaller contig require reads to be removed from the final BAM. In the latter case, a list of reads to be deleted is maintained, which correspond to reads covering the deleted region in the original BAM. BAMSurgeon requires approximately 4GB of memory per thread if using the Burrows-Wheeler Aligner (BWA). Its runtime varies depending on the number, variety and locations of the mutations, as well as the depth of the original BAM. On average, runtime is about 2-3 minutes per SV per thread followed by several hours to integrate all mutations into the output BAM, for a deeply sequenced (*e.g.* 60x) genome. These are wallclock times, with the majority being spent in writing reads into the BAM file.

**Fig. 1.**
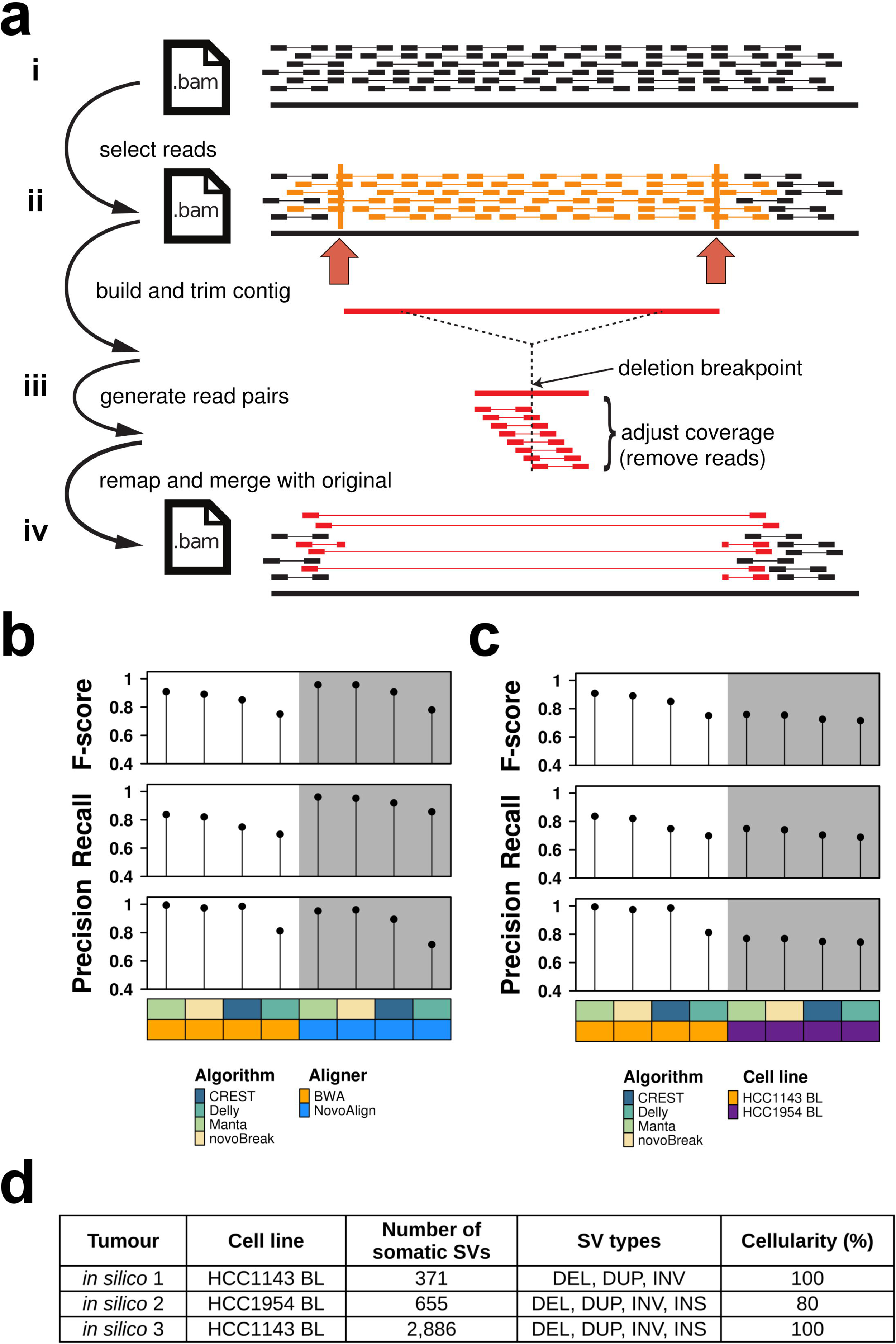
| BAMSurgeon simulates SVs in genome sequences. Method for adding SVs to existing BAM alignments. **a** Overview of SV (*e.g.* deletion) spike-in: Starting with an original BAM (i), a region (ii) is selected where a deletion is desired. iii) Contigs are assembled from reads in the selected region, and the contig is rearranged by deleting the middle. The amount of contig deleted is a user-definable parameter. Read coverage is generated over the contig using wgsim to match the number of reads per base in the original BAM. Since the deletion contig is shorter than the original, fewer reads will be required to achieve the equivalent coverage. iv) Generated read pairs include discordant pairs (*i.e.* paired reads that do not align to the reference genome with the expected relative orientation and inner distance) spanning the deletion and clipped reads (*i.e.* reads that are only partially aligned to the reference). Reads mapping to the deleted region of the contig are not included in the final BAM. **b,c** To test the robustness of BAMSurgeon with respect to changes in (**b**) aligner and (**c**) cell line, we compared the ranks of CREST, Delly, Manta and novoBreak on two new tumour-normal data sets: one with an alternative aligner, NovoAlign, and the other on an alternative cell line, HCC1954 BL. Callers were scored with *f* = 100 bp (Additional file 1: Figure S2b); Manta retained the top position, independent of aligner and cell line. **d** Summary of the three *in silico* (IS) tumours described here. Abbreviations: DEL, deletion; DUP, duplication; INV, inversion; INS, insertion.

### Validation of simulated somatic SVs

To validate SVs simulated by BAMSurgeon, we performed a series of quality-control experiments analogous to those performed to validate simulated SNVs [10]. Briefly, we used BAMSurgeon to generate synthetic tumour-normal pairs, with the same set of target mutations, that differ by the division of reads into tumour and normal sequence sets, aligner or cell line. The target mutation set was designed to generate a synthetic tumour with a baseline level of complexity and thus did not include insertions. We ran four SV callers using default parameters on each pair: two widely used callers, CREST [16] and Delly [9], and two callers developed over the course of the Challenge, Manta [17] and novoBreak [18]. We did not optimize parameters for the callers since the goal of this validation was not to identify the best caller, but instead to verify that the caller ranking is maintained across analogous datasets.

The definition of a SV suggests different scoring schemes for measuring the performance of a caller. All SVs can be defined by at least one breakpoint; deletions, duplications and inversions are SVs defined by a pair of breakpoints that in turn defines a genomic region. Thus, we compared called SVs to gold-standard SVs based on i) region overlap or ii) breakpoint closeness (Table 1, Additional file 1: Figure S2). The Challenge initially used a scoring scheme based on region overlap (at least one or more bases in common; Additional file 1: Figure S2a). Here we focus on the breakpoint closeness scheme since it is well suited for all types of SVs. A called SV that is sufficiently similar to a known SV based on such criteria was considered a true positive; otherwise, a false positive. We used such annotations to assess the performance of a caller in terms of precision (fraction of calls that are true), recall (fraction of known SVs called) and *F*-score (harmonic mean of precision and recall).

**Table 1.** | Caller scoring schemes.

| **Basis of comparison** | Region Overlap<br>(Additional file 1: Figure S2a) | Breakpoint Closeness<br>(Additional file 1: Figure S2b) |
| --- | --- | --- |
| **Description** | SVs match if there is sufficient overlap, determined with a Jaccard threshold parameter, between the genomic region associated with the called SV and that of the known SV | SVs match if the breakpoints of the called SV are sufficiently close to the those of the known SV, *i.e.* breakpoints are within $f$ bp of one another where $f$ is a flank parameter |
| **Strengths** | identifies genomic regions affected by the known SVs | suited to all types of SVs<br>evaluates precision of breakpoint predictions, facilitating subsequent breakpoint validation |
| **Weaknesses** | some SVs are not accurately defined by genomic regions, e.g. an insertion may be characterized by a single breakpoint<br>need criteria to define sufficient overlap | need criteria to define sufficient closeness |

We performed several quality-control experiments. First, the caller ranking (by *F*-score) was independent of the random division of reads: Manta > novoBreak > CREST > Delly (Additional file 1: Figure S3a,b). Second, the same ranking was observed when alignments were generated either using the BWA or NovoAlign with and without INDEL realignment (*i.e.* local realignment to minimize mismatches across reads due to INDELs relative to the reference genome), indicating that the ranking was independent of the aligner used (Fig. 1b, Additional file 1: Figure S3c). Lastly, when the genomic background was varied by using HCC1143 BL or HCC1954 BL sequence data, the caller ranking was largely independent of the cell line: Manta and novoBreak retained first and second place, respectively, while CREST and Delly swapped places, although their *F*-scores were very similar to each other when HCC1954 BL was used (Fig. 1c, Additional file 1: Figure S3d). Overall, these results show that simulated SVs are robust to changes in the read division, aligner and genomic background.

### Crowd-sourced benchmarking of somatic SV calling

The SV component of the Challenge consisted of the same three synthetic tumour-normal data sets used in the SNV component [10]. Briefly, the data sets were derived from existing cell line sequence data (thus minimizing data access restrictions) and *in silico* tumours 1-3 (IS1-IS3) were generated with increasing complexity (Fig. 1d). In terms of SVs, breakpoint locations were randomly selected and the tumours had increasing mutation rates (371 *vs.* 2,886 somatic SVs in IS1 and IS3, respectively). Moreover, IS1 contained deletions, duplications and inversions while IS2 and IS3 additionally contained insertions. Like the SNV component, the SV component of the Challenge was implemented using the Dialogue for Reverse Engineering Assessments and Methods (DREAM) framework. Briefly, information about the Challenge was shared on its website [19], participants registered online, downloaded a data set, applied their SV calling pipelines to the data set and submitted the results in Variant Call Format (VCF) v4.1. IS1-IS3 were released sequentially, each data set had its own competition phase and participants could make multiple submissions for each data set. Each tumour genome was divided into a training set and a testing set by holding out a portion of the genome. During the competition phase, leaderboards showed performance measures on the training set. After the competition closed, leaderboards also showed performance measures on the whole genome (training + testing sets).

The Challenge administration team prepopulated the leaderboards with two submissions and the community provided 204 submissions from 15 teams (Table 2, Additional file 2). A list of all submissions, and descriptions of pipelines used to generate them, can be found in Additional files 3 and 4, respectively. The submissions were surprisingly discordant in format. For example, between 5.5-11% of all submissions specified SV types that are not in the VCF controlled vocabulary for types (Additional file 5). For this reason, and the ambiguity of specifying SV types (*i.e.* the same SV can be specified with a specific type, or by specifying the breakpoints and break-end adjacencies), type specifications were ignored when scoring submissions. Team ranking varied with the stringency of the scoring (Additional file 1: Figure S2d-i). For simplicity, we focused on scoring with *f* = 100 bp due to the balance between the median and variance of the resulting *F*-scores (Additional file 1: Figure S4). While the top-performing teams achieved near maximal precision on the simplest tumour, IS1, their recall remained less than 0.9 (Fig. 2a), and decreased further on the other tumours (Additional file 1: Figure S5a,b). On all three tumours, *F-* scores on the training and testing sets were highly correlated (Spearman’s rank correlation coefficient (ρ) ≥ 0.98; Fig. 2b, Additional file 1: Figure S5c,d). However, the slightly elevated *F-* scores in the training sets observed for IS1 and IS2 may reflect minor overfitting; overfitting occurs when a statistical model is tuned to the training set, limiting generalizability. Notably, the total number of somatic SV mutations in IS3 is >4x that for IS1 and IS2 (Fig. 1d). Conversely, the percentage of mutations used for training is greater for IS1 (93%) and IS2 (92%) *vs.* IS3 (89%). Sampling from the IS3 mutations, we simulated training and testing sets of different sizes, and computed the differences between the *F-* scores on the training sets and the *F*-scores on the testing sets. We found that that the differences tend to be greater when the percentage of mutations used for training is greater (Additional file 1: Figure S5e). This suggests that the *F-* score differences observed for IS1 and IS2 are at least in part an artefact of training set size.

**Fig. 2.**
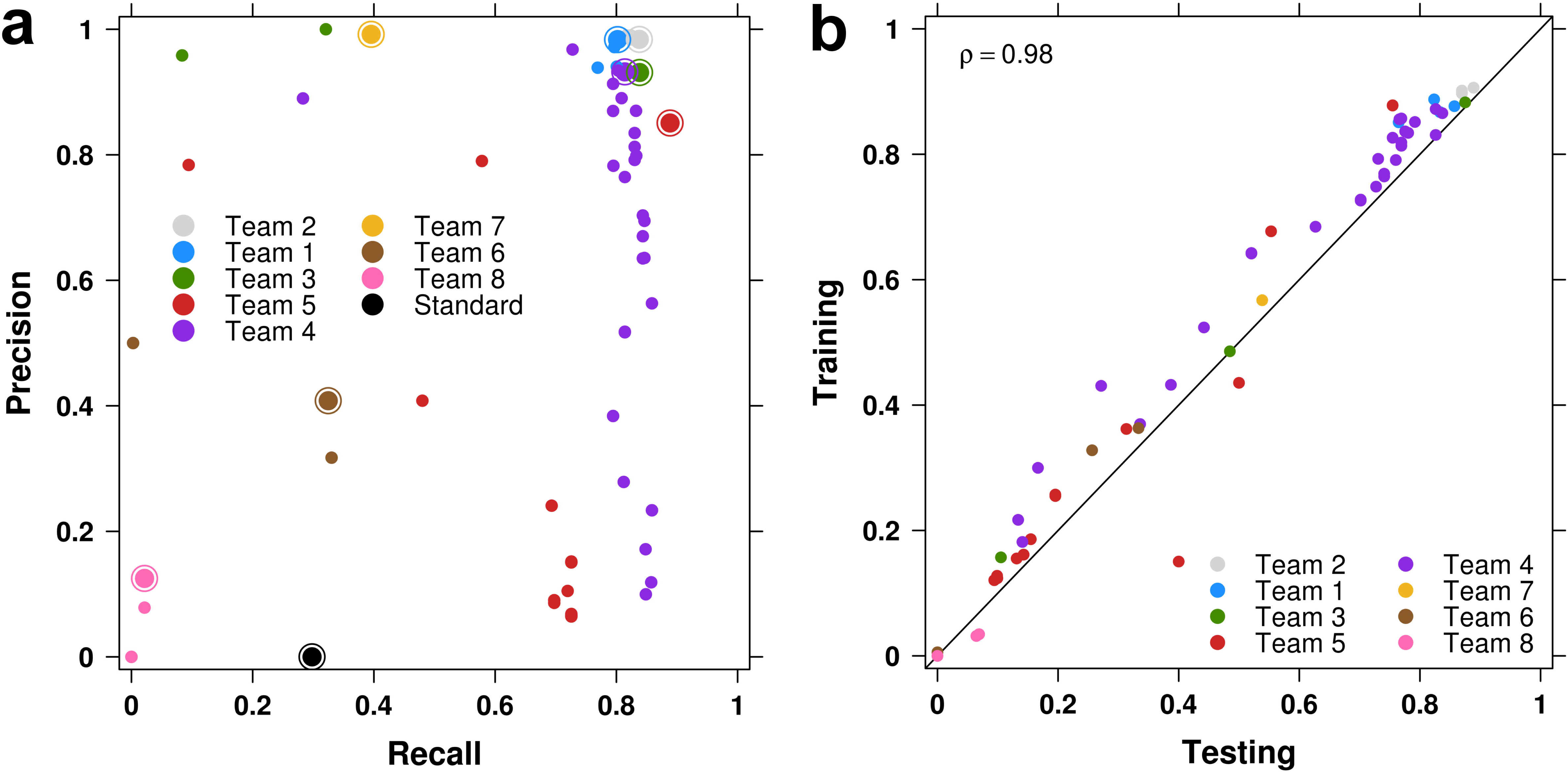
| Overview of the SV Calling Challenge submissions. a Precision-recall plot of IS1 submissions. Each point represents a submission, each colour represent a team and the best submission from each team (top *F-* score) is circled. The “Standard” point corresponds to the reference point submission provided by Challenge organizers. **b** The *F-* scores of submissions on the training and testing sets are highly correlated for IS1 (Spearman’s ρ = 0.98), falling near the plotted *y* = *x* line.

**Table 2.** | Teams.

| Team ID | Method name | Institute |
| --- | --- | --- |
| Standard | BreakDancer | Challenge Administrators |
| Team 1* | Delly<sup>1</sup> | European Molecular Biology Laboratory (EMBL) |
| Team 2 | Manta | Illumina, Inc. |
| Team 3 | Meerkat | Harvard Medical School |
| Team 4 | novoBreak | MD Anderson Cancer Center |
| Team 5 | deStruct<sup>2</sup> | BC Cancer Agency Research Centre |
| Team 6 | SWAN | Wharton School |
| Team 7 | SmuFin | Barcelona Supercomputing Center |
| Team 8 | not available | Peking University |
| Team 9* | deStruct<sup>2</sup> | Simon Fraser University |
| Team 10 | GROM | Rutgers University |
| Team 11 | Pindel | McDonnell Genome Institute |
| Team 12 | PAVFinder | Canada's Michael Smith Genome Sciences Centre |
| Team 13 | Delly<sup>1</sup> | Georgetown University Medical Center,<br>National Cancer Institute |
| Team 14 | Subread | Walter and Eliza Hall Institute of Medical Research |
| Team 15 | CX | Wharton School |

Teams that made submissions for IS1, IS2 and/or IS3, the names of the SV detection methods they used and the institutes to which they belong.
<sup>1</sup>Delly was developed by Team 1 and also used by Team 13.
<sup>2</sup>deStruct was developed by Team 9 and also used by Team 5.

### Pipeline optimization

The within-team variability in *F* -scores accounts for 21-43% of the total per-tumour variance in *F* -scores. The large variability in submissions by certain teams highlights the impact of tuning parameters during the Challenge (Fig. 3a, Additional file 1: Figure S6a,b). In comparing the initial (“naive”) and best (“optimized”) submissions of each team, for each tumour, the maximum *F-* score improvement was 0.75 (from 0.12 to 0.87 by Team 5 for IS1), and the median improvements were 0.20, 0.01, and 0.07 for IS1, IS2 and IS3, respectively (Fig. 3b). At least 33% of teams improved their *F-* score by at least 0.05 and at least 25% of teams improved it by more than 0.20, depending on the tumour. Despite these improvements by parameterization, team rankings were only moderately changed: if a team’s naive submission ranked in the top three, their optimized submission remained in the top three 66% of the time (Fig. 3c).

**Fig. 3.**
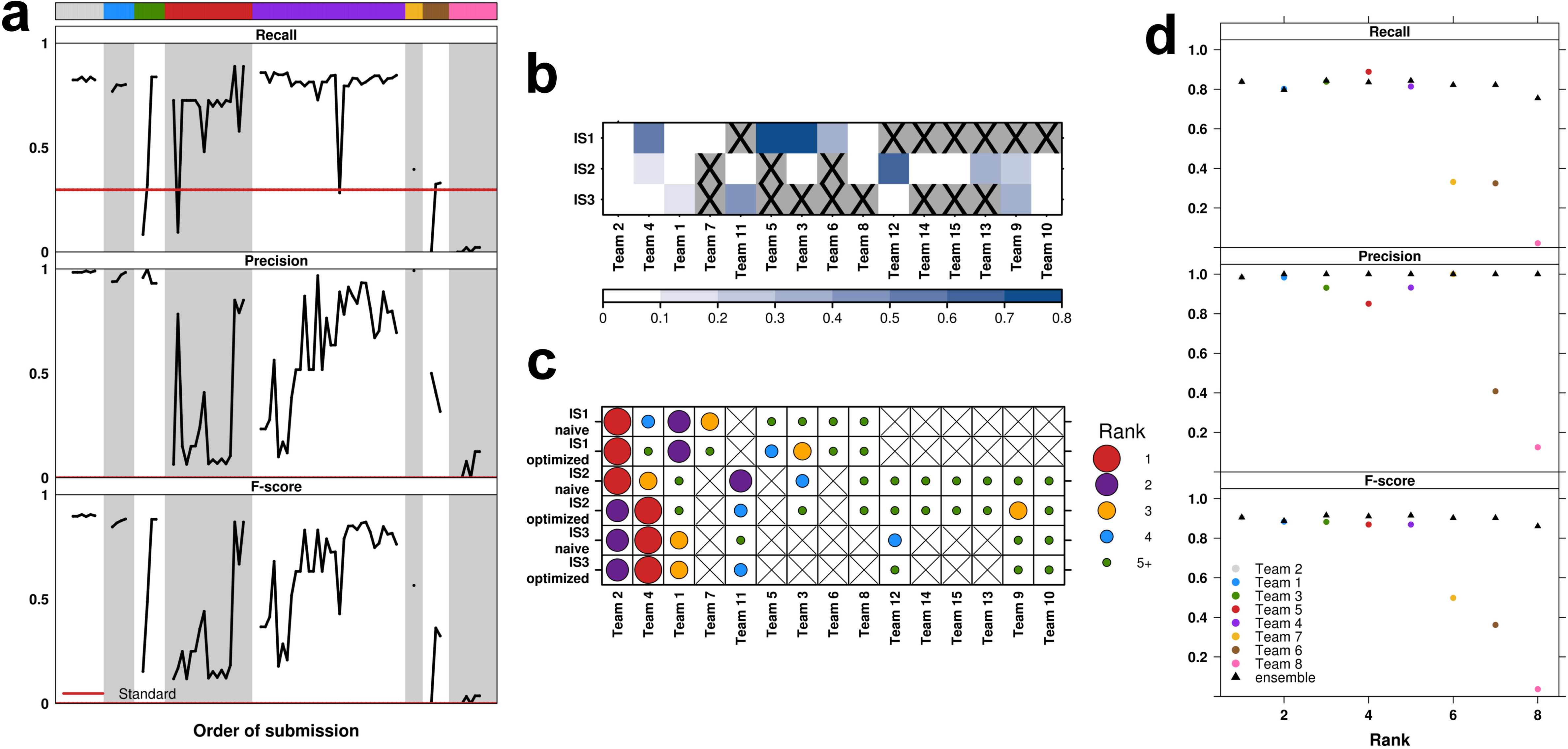
| Performance optimization by parameterization and ensembles. a Recall, precision and *F-* score of all IS1 submissions plotted by team, then submission order. Teams were ranked by the *F*-score of their best submission, colour coding (top bar) as in Fig. 2. The “Standard’” lines correspond to the reference point submission provided by Challenge organizers. **b** For each tumour, the improvement in *F-* score from the initial (“naive”) to the best (“optimized”) submissions of each team. Darker shades of blue indicate greater improvement. **c** For each tumour, team rankings based on their naive or optimized submissions. Larger dot sizes indicate better ranks by *F*-score. **b,c** An “X” indicates that the team did not make a submission for the specific tumour (or changed team name). **d** Recall, precision and *F-* score of ensembles versus individual submissions for IS1. At the *k* th rank, the triangles indicate performance of the ensemble of the top *k* submissions, and the circles indicate performance of the *k* th ranked submission. The ensemble analysis focused on the best submission from each team.

Given the crowd-sourced nature of the Challenge, we explored “wisdom of the crowds” as an approach to optimize performance [20,21]. Specifically, we aggregated SV calls into an ensemble by first identifying sufficiently similar calls in the majority of the top *k* submissions. Pairwise distances between calls from different submissions were computed (*i.e*. a breakpoint-length distance that incorporates distances between breakpoints and differences in SV length, Additional file 1: Figure S2c), and those calls with distances less than a selected threshold (equal to *f*, for consistency) were considered to represent an equivalent called SV event. The chromosome together with the median start and end positions of a set of similar calls would then define a single ensemble SV prediction. We considered two variations of this approach: i) a baseline approach with ensembles of the best submission from each team, and ii) a conservative approach with ensembles of all submissions (where the top *k* may include multiple submissions from the same team) and more stringent aggregation of called SVs (see Methods). The baseline ensembles were found to have *F-* scores comparable to that of the best submission (*e.g.* for IS1, the best ensemble and submission have *F*-scores of 0.92 and 0.91, respectively; Fig. 3d, Additional file 1: Figure S7b). However, the ensembles had lower *F*-scores than the best submission for IS2 (Additional file 1: Figure S7a). When *k* > 15, we found that the conservative ensemble *F*-scores drop further below that of the best submission (Additional file 1: Figure S7c-e; *e.g.* for IS1, the best ensemble with *k* > 15 and the best submission have *F*-scores of 0.83 and 0.91, respectively); these ensembles incorporate submissions from the top three teams, at least. In contrast, the precision of all ensembles (range: 0.993-1.00) was similar or slightly improved compared to that of the best submission. Thus, any changes in the ensemble *F-* scores were mostly influenced by the changes in recall as *k* varied.

### Error characteristics

We next exploited the large number of independent analyses to identify characteristics associated with false negatives (FNs) and false positives (FPs). For example, error profiles differed significantly between subclonal populations in IS3, with greater FN rates for mutations present at lower VAFs (Additional file 1: Figure S8; one-sided Wilcoxon signed rank *P* = 0.02 for VAF = 0.2 vs. 0.33, *P* = 0.04 for VAF = 0.33 vs. 0.5, *n* = 7). We also selected the best submission from each team (by *F*-score) and focused on 14 variables associated with breakpoint positions, representing factors like coverage and mapping quality (Additional file 6). Several of these variables were associated with false-positive rates; in particular, tumour coverage (*R* > 0.24), bridging reads count (the number of reads that bridge a putative breakpoint, *R* > 0.21) and mapping quality (*R* < -0.29), have stronger associations with FPs for both IS2 and IS3, compared to other variables (Additional file 1: Figure S9a, S10-S25). By contrast, few were associated directly with false-negative rates (0 ≤ |*R|* ≤ 0.15; Additional file 1: Figure S9b, S10-S25).

To evaluate whether these variables, and additional categorical variables, contribute simultaneously to somatic SV prediction error, we generated two Random Forests (non-parametric learning models that can trivially merge multiple data types) [22] for each team to assess variable importance for FN and FP breakpoints separately. FN breakpoints are associated with variables such as high bridging reads count and strand bias (Fig. 4a,c,e,g,i; Additional file 1: Figure S26a). FP breakpoints are generally associated with variables such as low mapping quality (Fig. 4b,d,f,h,j; Additional file 1: Figure S26b).

**Fig. 4.**
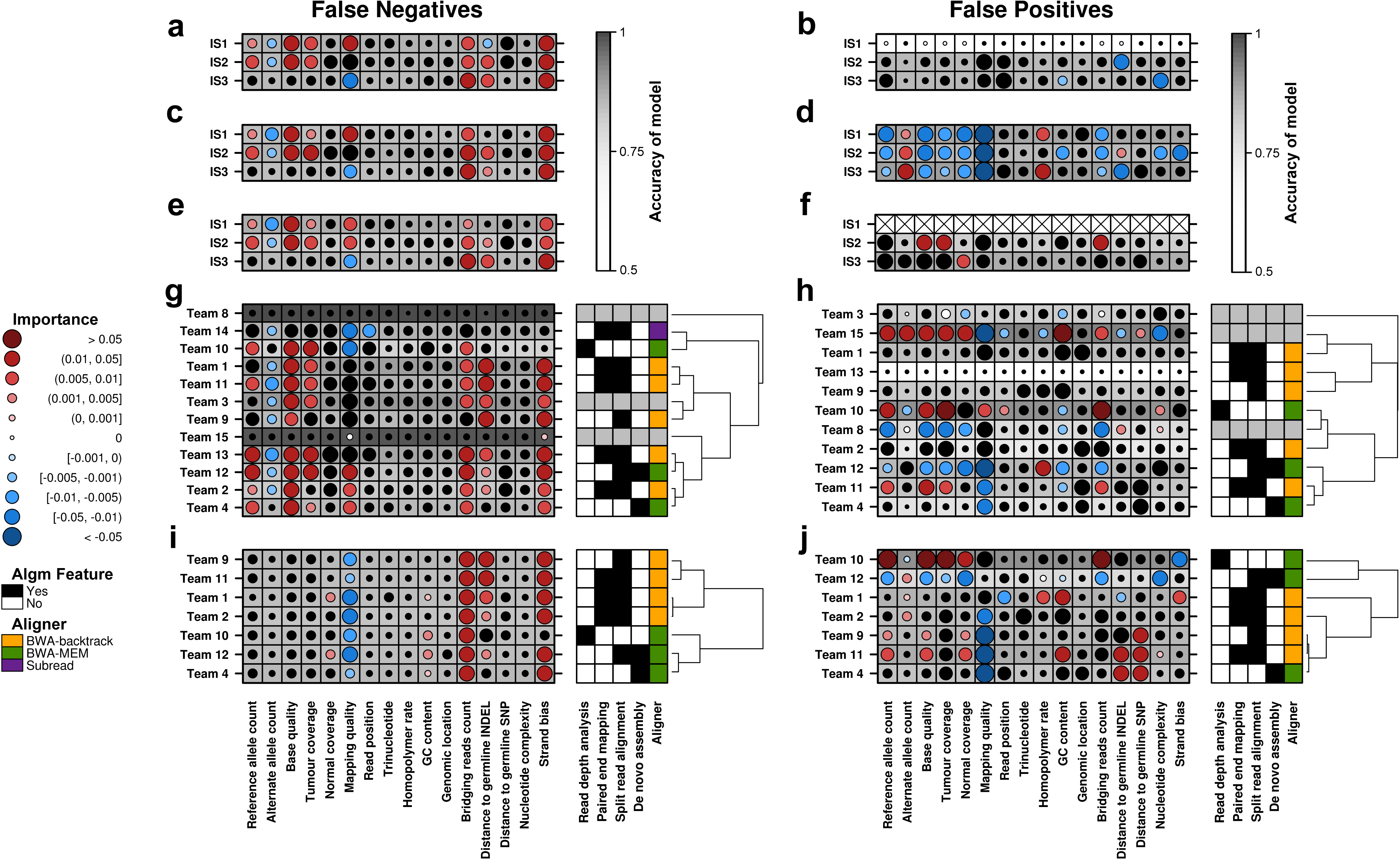
| Characteristics of prediction errors. Random Forests assess the importance of 16 sequence-based variables for each caller’s FN (**a,c,e,g,i**) and FP (**b,d,f,h,j**) breakpoints. Each panel shows variable importance on the left, where each row represents the best performing set of predictions by the given team/caller (on the given *in silico* tumour), and each column represents the indicated variable. Dot size reflects variable importance, *i.e.* the mean change in accuracy caused by removing the variable from the model (generated to predict erroneous breakpoints). Colour reflects the directional effect of each variable (red and blue for greater and lower variable values, respectively, associated with erroneous breakpoints; black for categorical variables or insignificant directional associations, two-sided Mann-Whitney *P* > 0.01). Background shading indicates the accuracy of the model (see colour bar). Variable importance for FN and FP breakpoints in each of the three tumours is shown for the following SV callers: CREST (**a,b**), Delly (**c,d**) and Manta (**e,f**). Manta only called two FPs in IS1; thus, variable importance for FP breakpoints could not be computed (indicated by Xs in the plot). Variable importance for FN and FP breakpoints in IS2 (**g,h**) and IS3 (**i,j**) is shown for each team. In the right plot (**g-j**), the first four columns indicate usage of the indicated algorithmic approaches by each team, and the last column indicates the aligner used. Grey indicates that algorithmic approaches and aligner are unknown for the given team. Abbreviations: Algm, algorithm; SNP, single-nucleotide polymorphism; INDEL, short insertion or deletion.

By executing specific SV callers, CREST (Fig. 4a,b), Delly (Fig. 4c,d) and Manta (Fig. 4e,f), with the same parameters on all three tumours, we identified tumour-specific error profiles. For example, the distance to the nearest germline INDEL tends to have stronger associations with errors in IS2 and IS3 compared to IS1 (Fig. 4a-e). Team-specific error profiles are more apparent with the FP breakpoint analysis. For example, Teams 8 and 10 have distinct FP profiles for the same tumour, IS2 (Fig. 4h); FPs by Teams 8 and 10 are negatively and positively associated with tumour coverage, respectively. Algorithmic approaches to SV calling from sequencing data include i) read depth analysis, ii) paired end mapping, iii) split read alignment, and iv) *de novo* assembly [23]. Some teams submitted sufficient algorithm details to determine the general approaches used, as well as the choice of aligner (Fig. 4g-j). Based on the available annotations, teams using the same aligner do not have error profiles that tightly cluster for all three tumours, suggesting that the aligner is not as strong a driver of those profiles, compared to the caller algorithm.

## Discussion

Crowd-sourced benchmarking challenges are ideal for questions where significant diversity in algorithmic approaches exists, particularly where individual methods are highly parameterized or computationally intensive [24,25]. The detection of variants from high-throughput sequencing data fits these criteria well: dozens of algorithms are in common use, many with complicated sets of parameters to tune and most requiring tens to hundreds of CPU hours to execute. We have quantified the critical importance of parameterization: it accounts for 21-43% of the variability in performance across the 204 submissions evaluated. This is comparable to the 26% of variability observed in somatic SNV detection benchmarking [10], and highlights the need for algorithm developers to continue to optimize parameters, provide guidance for their tuning and work toward automating their selection to make their software easier to use.

Scoring SV detection is complicated by the diversity of SVs. While some SV types may be well-characterized by overlap-based scoring methods, others benefit more from breakpoint-based scoring, and the choice of scoring metric and stringency parameters must be tuned to specific biological questions of interest. For example, breakpoint identification is critical when considering translocations -- especially those generating candidate fusion proteins -- while overlap of the called and known regions is much more important for copy-number analyses. Moreover, it may be useful to adapt scoring (*e.g.* by using a range of stringency parameter values) to identify SVs in certain contexts (*e.g.* with breakpoints in repetitive regions) that are still detectable by given tools, but with less precision. Taken together, SV diversity is an important consideration for the development of standards for scoring SV detection.

The “wisdom of the crowds” is the idea that an ensemble of multiple algorithms can significantly outperform any individual method. Several crowd-sourced benchmarking competitions from diverse fields have shown great success in combining submissions from contestants to achieve a high-performing meta-predictor including challenges for somatic SNV detection [10], gene regulatory network inference [21] and mRNA-based prognostic signatures for breast cancer [20]. By contrast, in somatic SV detection, we do not have clear evidence that an ensemble improves on the best individual method consistently across different tumours. Specifically, the majority vote approach works very well for somatic SNV detection, yet it appears to fail for somatic SV detection. This may reflect the large diversity in the biases of each individual algorithm (Fig. 4). Rather than focus on commonalities through a majority vote, it may be more beneficial to leverage the strengths of individual algorithms. This might be achieved by using machine learning to optimize the weighting of the algorithms for specific input patterns. For example, an aggregating classifier could learn, if there is a sizable difference in coverage in the tumour versus normal samples near given candidate breakpoints, the calling algorithms that use read depth analysis should have more weight. The overall approach could involve the following general steps: 1) apply all algorithms of interest to a given tumour-normal dataset and take the union of all resulting call sets to define a list of candidate SVs; then for each candidate, 2) compute sequence features (*e.g.* coverage) around the candidate breakpoints, and 3) provide computed features and confidence scores from individual algorithms as input to an aggregating classifier that will indicate whether or not the candidate is likely to be a true SV. In fact, a similar approach is behind the SMC-DNA Meta-pipeline Challenge [26] for benchmarking pipelines that aggregate calls from different SNV detection algorithms. In practise, analogous efforts for SV detection would require additional considerations such as the identification of i) an optimal method for merging similar yet different calls (due to imprecise breakpoint calling) when compiling the list of candidate SVs, ii) the most informative sequence features for guiding the relative weighting of individual algorithms (*e.g.* variables in Figure 4), and iii) an optimal scoring method (as mentioned above). Thus there is a need for continued development of new, more complex ways to integrate multiple somatic SV detection methods [27].

Given the paucity of gold-standard benchmarking data for somatic SVs, the creation of the simulated datasets and the associated leaderboards constitutes a major contribution of this Challenge. Ideally, a simulated dataset depicts realistic mutations through realistic sequence reads. The synthetic tumours generated for the Challenge only represent straightforward SV types (duplication, deletion, insertion, inversion) and cover relatively small regions. Subsequent enhancements to BAMSurgeon have added support for additional SV types including translocations and complex SV combinations, enabling simulations to more completely capture the complexity of tumour genomes and by extension, challenge SV callers in different ways. For each SV, simulated reads are generated (*via* wgsim) from a re-arranged contig, where the original contig is constructed from real reads. Despite the basis on real reads, the simulated reads do not necessarily reflect the non-uniform coverage that may arise during preparation of real samples, for example [28]. There are other read simulators that learn biased-coverage trends from real data and use them to generate reads (*e.g.* [29]) that could be used by BAMSurgeon; however, it is an on-going challenge to simulate biases of real sequencing data as sample preparation methods and sequencing technologies vary and/or advance. In fact, one could sequence the same ‘normal’ sample twice to capture inter-sample variability, with one replicate converted into a synthetic tumour sample using BAMSurgeon. Nevertheless, there are distinct advantages to benchmarking on simulated datasets. It is dramatically easier to simulate large numbers of tumours, or to create tumours with highly divergent mutational properties, leading to well-supported estimates of per-tumour caller accuracy. This enables our strategy of generating synthetic tumours of increasing complexity (*e.g.* with other SV types and/or haplotype structure by using phased sequence data) whereby the impact of the complexity introduced at each step can be assessed. With the three synthetic tumours described here, we observed that caller ranking varied across tumours and we expect it to vary with a broad range of tumour characteristics including coverage, normal contamination, complexity of the SVs, the number of mutations adjacent to breakpoints and others, as they each present different challenges. It is possible to identify strengths and weaknesses of an individual caller by comparing its tumour-specific error profiles. Moreover, synthetic tumours can be designed to test the limits of callers. These advantages highlight the usefulness of synthetic datasets for benchmarking callers, and until synthetic datasets are completely realistic, they will serve as important complements to real datasets.

While 15 teams participated in the actual competitive phase of the Challenge, 8 teams have exploited the IS1-3 benchmarking resources since the competition, making 73 submissions to benchmark their methods for pipeline evaluation and development. Evaluations based on the first synthetic tumours, the simplest by design, provide lower-bounds on the error rates. As subsequent updates to BAMSurgeon enable the generation of more complex and realistic tumours, the corresponding error rates using these simulations will approach their upper-bounds. We hope that journals and developers will begin to plan for benchmarking on these standard datasets, including simulated ones, as a standard part of manuscripts reporting new somatic SV detection algorithms.

## Conclusions

Analysis of the error profiles of the Challenge submissions showed that somatic SV calling is a distinctly harder problem than somatic SNV calling even given a relatively simple set of SVs, with individual pipelines having complex and unique error profiles. Parameterization was a critical factor in determining the performance of teams. Finally, we demonstrate that, unlike almost every past DREAM Challenge, somatic SV prediction does *not* benefit from the “wisdom of the crowds” -- simple voting of multiple prediction pipelines does not yield improved predictions in this instance. The synthetic tumours and somatic SV detection leaderboards remain available as a community benchmarking resource.

## Methods

### Simulation of SVs by BAMSurgeon

SV support in BAMSurgeon has evolved throughout the Challenge, largely as a result of constructive feedback from participants. Our descriptions of BAMSurgeon’s method for simulating SVs is current as of commit (*i.e.*, version) b851573474 of the code available at [30]. As input, BAMSurgeon (addsv.py) requires an indexed reference genome, a pre-existing BAM file (Additional file 1: Figure S1a), and a list of intervals (Additional file 1: Figure S1b) along with the SV type and parameters (see manual [31]). The intervals should be wide enough that local sequence assembly is successful in generating a contig that spans at least 2x the expected library size in the input BAM file. Intervals for which a sufficiently long contig cannot be generated are rejected, where the exact definition of ’sufficiently long’ is an optional parameter. Note that it may be less likely to obtain long contigs from genomic regions that are more difficult to sequence, and by extension, less likely to simulate SVs in such regions. Intervals which contain too many discordant read pairs (again, potentially indicating regions that are difficult to sequence) are also rejected, subject to a parameter. Following local assembly, the contig is re-arranged: the specific rearrangements for each supported SV type are illustrated in Fig. 1a (step and Additional file 1: Figure S1c,e,g. The assembled contig is then re-aligned to the target interval in the reference genome (exonerate -- bestn 1 -m ungapped) and is trimmed based on the start and end coordinates of this alignment. Read pairs corresponding to trimmed contig sequence are removed from further consideration.

Read coverage is generated over the rearranged contig using a read simulator (wgsim -e 0 -R 0-r 0), to achieve the same average depth as the input BAM file, which has the effect of creating split reads relative to the reference genome supporting SV detection. For a deletion, the number of reads required to achieve (*e.g.*) 30x coverage is fewer than the number of reads required to reach 30x coverage prior to the deletion, so reads must be removed from the original BAM (Fig. 1a, step iv). Inversely, for duplications and insertions additional reads need to be added to the original BAM (Additional file 1: Figure S1d,h). Inversions generally do not affect coverage (Additional file 1: Figure S1f). To ensure any reads removed actually correspond to the deleted region of the contig, the locations of reads in the assembled contig are tracked. The number of reads to be replaced, added, or deleted is scaled with the desired allele fraction. Finally, any read pairs in the original BAM corresponding to reads altered in the simulated SV are replaced, any read pairs marked for deletion are removed from the original BAM, and any additional read pairs generated are added. It is recommended that the resulting altered BAM be post-processed (with postprocess.py) to ensure compliance with the SAM format specification (see manual [31]).

### Synthetic tumour generation

Synthetic tumours were prepared by partitioning high-coverage BAMs from ’normal’ cell lines into two groups of reads, picking read pairs at random as described previously [10]. Alternatively, one could sequence the same ‘normal’ sample twice to capture inter-sample variability, with one replicate converted into a synthetic tumour sample using BAMSurgeon. For the three *in silico* challenges, non-overlapping regions were selected at random for SV addition, resulting in 371 variants added for IS1, 655 for IS2, and 2,886 for IS3 (Fig. 1d). Variant input files are available in Additional file 7. SVs were added using addsv.py with assembly GRCh37/hg19 as the reference genome and default parameters except where noted. For IS3, to simulate subclones a file specifying CNV fractions over SV regions was input via option -c to specify the variant allele frequency (VAF) of the spiked-in variants at either 0.5, 0.33, or 0.2 (Additional file 7). The output BAMs were post-processed to account for any inconsistencies introduced due to remapping and merging of reads supporting SVs using the script postprocess.py included with BAMSurgeon. The BAMs were further adjusted with RealignerTargetCreator and IndelRealigner from the Genome Analysis Toolkit (v.2.4.9). All tumour-normal pairs generated via BAMSurgeon are verified for adherence to the SAM/BAM format specification using the ValidateSamFile tool included in the Picard tool set [32]. Truth VCF files, *i.e.* files specifying simulated mutations, for SVs were generated using the script etc/makevcf_sv.py and merged with truth files for SNP and INDEL locations, where applicable. SAMtools was used throughout to split, merge, sort, and index BAMs, and also index FASTA files. Details on the specific BAMSurgeon commits used for generating each tumour, as well as other tumour details are given at [33].

### Validation of BAMSurgeon

To validate BAMSurgeon’s ability to simulate somatic SVs, we compared the output of four algorithms -- two widely used SV callers, CREST [16] and Delly [9], and two callers developed over the course of the Challenge, Manta [17] and novoBreak [18] -- on the IS1 tumour-normal data set, and analogous datasets generated with the same spike-in set of mutations, but with an alternate aligner (NovoAlign v.3.00.05 [34]), cell line (HCC1954 BL) or read division. We did not optimize parameters for the callers since the goal of this validation was not to identify the best caller, but instead to verify that the caller ranking is maintained across analogous datasets.

Each tumour-normal pair was processed by CREST (v1.0) to extract soft clipping positions for each chromosome separately, using default parameters. This data was then used by CREST to call somatic SVs using the default protocol, and we converted the output into VCF v4.1 format. Somatic SVs were called from each tumour-normal pair using Delly (v0.5.5) with default parameters. Calling was performed on each chromosome separately for all supported SV types except for translocations, and we converted the translocation output into VCFv4.1 format. Calls with MAPQ < 20, PE < 5, or labeled as “LowQual” or “IMPRECISE” were filtered out. Somatic SVs were called from each tumour-normal pair using Manta (v0.26.3) with the following parameters: -m local -j 4 -g 10. Lastly, somatics SVs were called from each dataset using novoBreak (v1.04) with a modification to ensure that sequence windows around breakpoints never go beyond the start of the chromosome. All sets of SV calls were scored with *f* = 100 bp and *j* > 0, callers were ranked based on *F-* score for each tumour-normal pair, and rankings were compared across pairs (Fig. 1b,c and Additional file 1: Figure S3).

### Preprocessing VCF files

We preprocess VCF files to parse out the SV-relevant details (*e.g.* the END coordinate in the INFO value or from the length of the REF sequence; if the END coordinate cannot be determined from those values, it is set to the POS coordinate), remove SVs that did not pass filters (as indicated by the FILTER values) and ensure consistent formatting between files. To ensure consistent formatting in accordance with the VCFv4.1 specification [35] we:

1. Add row entries to ensure that each MATEID specification has a corresponding pair of entries, where only a single entry is provided
2. Re-assign IDs and MATEIDs to ensure unambiguous references to entries
3. Where possible, replace SVTYPE = BND entries with entries specifying SVTYPE = {CNV, DEL, DUP, INS, INV} in accordance with REF, ALT and EVENT values

Testing set SVs are indicated in the truth VCF file with the addition of masked genomic regions specified with CHROM, POS and END values indicating the chromosome, start and end coordinates, and SVTYPE = MSK. Specifically, a SV where ≥ 50% of the corresponding region overlaps a masked region is allocated to the testing set; otherwise, it is in the training set.

### Structural variant scoring

Our scoring approaches evaluate the accuracy of a set of called SVs and requires input VCF files specifying: i) called SVs, and ii) true/known SVs. Generally, a called SV that is sufficiently similar to a known SV based on specific criteria (Table 1) is considered a true positive (TP); otherwise, a false positive (FP). Also, a known SV that is sufficiently similar to a called SV is considered a TP; otherwise, a false negative (FN). Our scoring supports two ways of quantifying similarity:

A. **Region overlap.** The Jaccard coefficient (*j*) is computed from the lengths (in bp) of intersection and union regions (Additional file 1: Figure S2a).
B. **Breakpoint closeness.** The distance (Δ, in bp) between called and known breakpoints is computed (Additional file 1: Figure S2b). If Δ ≤ *f* (where *f* is a flank threshold parameter), a relative closeness is computed, *c’* = 1 -Δ/*f*. The overall closeness (*c*) is defined as the geometric mean of the *c’* values for the start and end breakpoints. If only one of the start and end breakpoints has Δ ≤ *f*, the called and known SVs are annotated as partially matching.

Unless otherwise specified, we scored with *f* = 100 bp. If there is an ambiguous matching of called SVs to known SVs by sufficient similarity, the similarity values (*j*/*c*) are used to identify an optimal one-to-one matching. First, we restrict the matching to the best match(es) for each called and known SV. If a SV has multiple best matches by similarity, we attempt to break the tie by favouring SVs with the same SVTYPE, and/or test/training set membership. If the best matching is still ambiguous, we then use corresponding similarity values together with the Hungarian algorithm to obtain a one-to-one matching (with the clue v0.3-48 R package [36]). Finally, SVs are annotated based on this matching. SVs that have sufficient similarity but are not in the final matching are annotated as partially matching. Mated breakpoints are initially annotated separately. If one is annotated as partially matching or as a TP, and the other is a FP, the FP annotation is replaced by a partial match annotation. Subsequently, each set of mated breakpoints is treated as a single SV.

These annotations are used to assess the performance of a SV caller in terms of precision = nTP/(nTP + nFP), recall = nTP/(nTP + nFN) and *F*-score (specifically, *F*_1_-score) = 2 x precision x recall/(precision + recall), where nTP, nFP and nFN represent the numbers of TPs, FPs and FNs, respectively. Partial matches are not counted in these computations. Unless otherwise specified, the precision, recall and *F*-score values presented here were computed on the testing and training sets combined. The best submission of a given team is defined as the team’s submission with the greatest *F*-score computed against all known SVs.

### Execution of challenge-based benchmarking

The SV component of the Challenge was executed concurrently with the SNV component, and the procedure has been described previously [10]. It was implemented using the Dialogue for Reverse Engineering Assessments and Methods (DREAM) framework. Briefly, information about the Challenge was shared on its website [19], participants registered online, downloaded a data set, applied their SV calling pipelines to the data set and submitted the results in VCFv4.1 format. IS1-IS3 were released sequentially, each data set had its own competition phase and participants could make multiple submissions for each data set. Each tumour genome was divided into a training set and a testing set. During the competition phase, leaderboards showed performance measures on the training set. After the competition closed, leaderboards also showed performance measures on the whole genome (training + testing sets), thus benchmarking the SV calling pipelines. The SV leaderboards for IS1 and IS2 were pre-populated with results from BreakDancer (v1.1.2_2013_03_08 [7]) run with default parameters; a reference point submission indicated labeled as “Standard” in figures and tables. Due to our exploration of multiple SV scoring methods in this manuscript, the leaderboard results are not completely consistent with the results presented here, but all raw and leaderboard data are available.

### Overfitting artefact analysis

Due to the order of magnitude greater number of SVs spiked into IS3, we simulated training and testing sets of different sizes by sampling from the IS3 training set. Specifically, we assessed mutation totals of 100 to 1000 (by increments of 100), and training sets that were 80-95% (by increments of 1%) of the total, by sampling each {mutation-total, training-set%} combination 100 times. For each sample, we computed *F_train_ -F_test_* for each IS3 submission where *F_train_* and *F_test_* are *F*-scores computed on the simulated training and testing sets, respectively. We then computed the median difference across samples to obtain a summary value for each submission, and finally show the median across submissions in Additional file 1: Figure S5e. (*F_train_* -*F_test_*) > 0 suggests overfitting but such values are an artefact of testing set size since no fitting/training was done in this analysis.

### Team variation

For each tumour-normal pair, we computed the percentage of variation in *F-* score, across all submissions, that is accounted for by within-team variation. Specifically, we computed the within-team sum of squares as a percentage of the total sum of squares.

### Definition of ensembles

We aggregated SV calls from *k* submissions into an ensemble set with the following general approach:

1. **BND filter.** Calls defined with SVTYPE = BND were excluded for simplicity.
2. **Compute call distances.** Pairwise distances (*d*, in bp) between remaining predictions were computed (*i.e*. a breakpoint-length distance that incorporates distances between breakpoints and differences in predicted SV length, Additional file 1: Figure S2c). Distances were only computed between predictions from different submissions.
3. **Generate sets of similar calls.** A distance less than a selected threshold (100 for consistency with *f*, see **Structural variant scoring**) indicated sufficiently similar calls. We assessed two variations:
  a. **Baseline.** We defined a graph such that vertices represented calls and edges connected sufficiently similar calls. We identified the connected components to define the sets of similar calls. Sets with median intra-set distances > *f* were refined. Specifically, the call with the greatest median distance to the other set members was iteratively removed until the median intra-set distance dropped below *f*, or the set became empty.
  b. **Conservative.** We used the added constraint that called SVs overlap by ≥ 1 bp to be treated as sufficiently similar. Sets of similar calls were constructed by iterating over the sufficient similarity pairs from least to most distant. If a pair did not contain a call in an existing call set, the pair was used to define a new call set. Otherwise, one call was already in a set, and the other was a candidate for addition to the same set via guilt-by-association. If the candidate came from a submission that was not already covered by the set, and its median distance to the existing set members ≤ *f*, it was added to the set. Any unprocessed pairs within or between the prediction sets at that stage were excluded from consideration.
4. **Majority vote filter.** Sets with calls from ≤ *k*/2 submissions were excluded; each remaining set covered the majority of submissions.
5. **Aggregate sets to define ensemble calls.** The chromosome together with the median start and end positions of each set of calls defined a single ensemble SV call.

An additional distinction between the baseline and conservative approaches is that the baseline approach only involved the best submission from each team whereas all submissions were used with the conservative approach. To investigate different ensembles of *N* submissions for the same tumour-normal pair, we first ordered the submissions by overall *F-* score, computed after excluding calls with SVTYPE = BND. We then generated an ensemble call set with the top *k* submissions, for *k* = 2.*N*. The performance of ensembles was compared to that of the individual submissions, after excluding calls with SVTYPE = BND (*e.g.* Fig. 3d).

### Error characterization

To characterize the errors made by a team, we assessed the team’s best submission for a given tumour-normal pair. We also assessed errors made by CREST, Delly and Manta when run, with the same protocols described in the **Validation of BAMSurgeon** section, on all three tumour-normal pairs. Characterizing FNs and FPs involved comparisons to TPs and true negatives (TNs), respectively. Moreover, we characterized errors at the level of breakpoints.

### Sampling true negatives

Given a set of submissions for the same tumour-normal pair, we identified the maximum number of FPs from a single submission, *m*. We then sampled ≥ *m* TNs for each submission, by sampling regions from the reference genome that satisfied these criteria:

1. length sampled from a log-normal distribution with mean and standard deviation equal to that of the logged lengths of the known SVs
2. start position is not in known gap and repeat regions
3. region does not overlap with any known SVs
4. region does not overlap with any SVs called in the submission

Some sampled regions qualified as TNs for multiple submissions. For IS2, we excluded Team 14’s submission because it had a very large number (17,806) of FPs, and thus was computationally problematic for the subsequent Random Forest analysis.

## Breakpoint annotations based on scoring

A single breakpoint may be associated with multiple (called/known) SVs, and therefore may be associated with multiple annotations depending on the scoring approach used, *i.e.* > 1 of {TP, FN, FP}. To remove ambiguity, we choose a single annotation for each breakpoint by prioritizing as follows: TP > FN > FP. This prioritization favours good performance (*i.e.* TP has highest priority) and then recall (*i.e.* FN > FP) since it appears to be a greater challenge than precision for SV calling (Fig. 2a, Additional file 1: Figure S5a,b). TN breakpoints should be unambiguous due to the way in which they were sampled (see above).

## Genomic variables

For each breakpoint position, we computed 16 genomic factors, 12 of which were previously described [10]. The additional genomic variables were computed as follows:

A. **Bridging reads count.** We used samtools v0.1.19 to identify reads in the tumour BAM mapped to a genomic region containing the window defined by the breakpoint position +/-1 bp. The bridging read count was defined as the number of identified reads. Note that a bridging read does not necessarily have a secondary mapping for part of the read, as one might expect for a split read.
B. **Distance to nearest germline INDEL.** Germline calls were obtained as previously described [10] and INDELs were parsed out. The distance of a breakpoint to the nearest germline INDEL was computed using BEDTools closest v2.18.2.
C. **Nucleotide complexity.** The sequence for the window defined by the breakpoint position +/-50 bp was extracted from the reference fasta file. The nucleotide complexity was defined as the entropy of the sequence: -S*p_x_* log_2_(*p_x_*) over *x* ∈ {A, G, C, T} where *p_x_* is the proportion of the sequence with *x* (case-insensitive).
D. **Strand bias.** We used samtools v0.1.19 to identify reads in the tumour BAM mapped to a genomic region containing the breakpoint position. The strand bias was defined as the proportion of these reads mapped to the + strand.

## Univariate analysis

To assess the relationship between each non-categorical variable and prediction error rates, we computed the Pearson correlation coefficient between the variable values and the proportion of teams with a FN/FP at the breakpoints, as well as the corresponding *P* value. Reference and alternative allele counts, base quality, tumour and normal coverages, bridging reads counts and distances to the germline SNP and INDEL were logged (base 10) prior to computing correlations (zeros were replaced with -1 instead of logged). For the categorical variables, trinucleotide and genomic location, the *P* value measured the significance of the variable in a fitted binomial model predicting the FN/FP rate at a breakpoint. A binomial model was fitted because it is a relatively simple model (and thus less prone to overfitting) to test the relationship between a categorical variable and a proportion variable (*i.e*. an error rate).

## Multivariate analysis

Random Forests were generated as previously described [10] with a few alterations. Here, a total of 16 genomic variables (Fig. 4) were used to build: i) a classifier to distinguish FN and TP breakpoints, and ii) a classifier to distinguish FP and TN breakpoints. A FP classifier was not generated for Team 7 with respect to IS1 since the team produced only one FP, and thus there was insufficient data to generate an accurate model. Conversely, a FP classifier was not generated for Team 14 with respect to IS2 since the team produced a very large number of FPs (17,806) that caused a failure to converge. Computation of the directional effect of variables was also as previously described [10].

Non-parametric tests (*i.e.* Wilcoxon and Mann-Whitney tests) were used throughout to avoid assumptions about the distributions of the tested populations; all tested populations had *n* ≥ 7.

The BEDTools suite (v2.18.2 [37]) was used with the bedR R package (v0.5.3 [38]) throughout. Plots were generated with the BPG (v5.3.9), lattice (v0.20-33) and latticeExtra (v0.6-26) R packages and R (v3.2.1) was used throughout.

## Declarations

### Availability of data and materials

Sequences files are available at the Sequence Read Archive (SRA) under accession number SRP042948. BAMSurgeon is available at Zenodo [39] and the code repository is available at GitHub [30]. Submission (Synapse IDs syn12628575, syn12628576 and syn12628577 for IS1-IS3, respectively) and known mutation (*i.e*. ground truth; Synapse IDs syn2354306, syn2399959 and syn2485207 for IS1-IS3, respectively) VCF files are available from the Challenge website [19] following registration and subsequent login at Synapse.

## Acknowledgments

The authors thank S.P. Shah, R.D. Morin and P.T. Spellman for helpful suggestions, L.E. Heisler and B.F. Huang as well as all the members of the Boutros lab for insightful discussions and technical support. The authors thank Google Inc. (in particular N. Deflaux) and Annai Biosystems (in particular D. Maltbie and F. De La Vega) for their ongoing support of the ICGC-TCGA DREAM Somatic Mutation Calling Challenge.

The ICGC-TCGA DREAM Somatic Mutation Calling Challenge Participants are: Bret D. Barnes, Inanc Birol, Xiaoyu Chen, Readman Chiu, Anthony J. Cox, Li Ding, Markus H-Y. Fritz, Andrey Grigoriev, Faraz Hach, Joseph K. Kawash, Jan O. Korbel, Semyon Kruglyak, Yang Liao, Andrew McPherson, Ka M. Nip, Tobias Rausch, S. Cenk Sahinalp, Iman Sarrafi, Christopher T. Saunders, Ole Schulz-Trieglaff, Richard Shaw, Wei Shi, Sean D. Smith, Lei Song, Difei Wang, Kai Ye.

## Funding

This study was conducted with the support of the Ontario Institute for Cancer Research to P.C.B. through funding provided by the Government of Ontario. This work was supported by Prostate Cancer Canada and is proudly funded by the Movember Foundation -Grant #RS2014-01. This study was conducted with the support of Movember funds through Prostate Cancer Canada and with the additional support of the Ontario Institute for Cancer Research, funded by the Government of Ontario. This project was supported by Genome Canada through a Large-Scale Applied Project contract to P.C.B., S.P. Shah and R.D. Morin. This work was supported by the Discovery Frontiers: Advancing Big Data Science in Genomics Research program, which is jointly funded by the Natural Sciences and Engineering Research Council (NSERC) of Canada, the Canadian Institutes of Health Research (CIHR), Genome Canada, and the Canada Foundation for Innovation (CFI). P.C.B. was supported by a Terry Fox Research Institute New Investigator Award and a CIHR New Investigator Award. K.E.H. was supported by a CIHR Computational Biology Undergraduate Summer Student Health Research Award. A.D.E was supported by an Australian Research Council Discovery Early Career Researcher Award DE150101117 and by the Mater Foundation. The following National Institutes of Health (NIH) grants supported this work: R01-CA180778 (J.M.S.), and U24-CA143858 (J.M.S.). The funders played no role in study design, data collection, data analysis, data interpretation or in writing of this manuscript.

## Author’s contributions

A.A.M., J.M.S and P.C.B. initiated the project. A.D.E. created BAMSurgeon. A.D.E, K.E., Y.H., K.E.H., J.C.B., M.R.K., T.C.N., S.H.F., G.S., A.A.M., J.M.S. and P.C.B. created the ICGC-TCGA DREAM Somatic Mutation Calling Challenge. A.Y.L., A.D.E., Y.H., K.E.H., S.M.G.E., V.H., K.D., Z.C., C.C., and T.N.Y. created datasets and analyzed sequencing data. A.Y.L., Y.H., K.E.H, and P.C.B. were responsible for statistical modelling. Research was supervised by K.C., S.H.F., J.G., G.S., D.H., A.A.M., J.M.S. and P.C.B. The first draft of the manuscript was written by A.Y.L. and P.C.B., extensively edited by A.D.E., K.E., A.A.M. and J.M.S. and approved by all authors.

## Ethics approval and consent to participate

Not applicable.

## Consent for publication

Not applicable.

## Competing interests

All authors declare that they have no competing interests.

## Additional files

Additional file 1: Figures S1-S26. (PDF 3.5 MB)

Additional file 2: Table S1. Challenge participation. (XLS 5 KB)

Additional file 3: Table S2. All competition-phase submissions evaluated with *f* = 100 and *j* > 0. (XLS 43 KB)

Additional file 4: Descriptions of pipelines used to generate submissions. (PDF 3.4 MB)

Additional file 5: Table S3. Invalid SV types. (XLS 8 KB)

Additional file 6: Table S4. Univariate error analysis. (XLS 14 KB)

Additional file 7: BAMSurgeon input files used to generate the three *in silico* tumour-normal pairs (IS1-IS3). (TAR.GZ 122 KB)

